# Brooklyn plots to identify co-expression dysregulation in single cell sequencing

**DOI:** 10.1101/2023.06.21.545951

**Authors:** Arun H. Patil, Marc K Halushka

## Abstract

Altered open chromatin regions, impacting gene expression, is a feature of some human disorders. We discovered it is possible to detect global changes in genomically-related gene co-expression within single cell RNA sequencing (scRNA-seq) data. We built a software package to generate and test non-randomness using ‘Brooklyn plots’ to identify the percent of genes significantly co-expressed from the same chromosome in ∼10MB intervals across the genome. These plots establish an expected low baseline of co-expression in scRNA-seq from most cell types, but, as seen in dilated cardiomyopathy cardiomyocytes, altered patterns of open chromatin appear. These may relate to larger regions of transcriptional bursting, observable in single cell, but not bulk datasets.

## INTRODUCTION

Single cell and single nucleus RNA-sequencing (scRNA-seq and snRNA-seq) have revolutionized our understanding of cell types and cell states, allowing deeper understanding of gene expression patterns beyond bulk sequencing of tissues(1). scRNA-seq analysis software packages, such as Seurat, cluster cells and assign them to unique identities, which can be visualized through multidimensionality reduction(2). These tools utilize the widely variable, and often unique, expression patterns of cells to form independent clusters of specific cell types or cell states. Within a defined cell cluster, it is known that absolute gene expression differences exist between individual cells. These can be the result of technical differences, such as depth of sequencing, cell health, or stochastic ambient RNA capture differences(3). Biological activity differences, such as cell cycle states, can also be identified through single cell approaches(4). We hypothesized that the cells within a specific cell type cluster may also vary by changes in their open chromosome domains, particularly in disease states. These areas likely coincide with enlarged (by chromosomal region) transcriptional bursts. Transcriptional burst locations would be stochastic and variable between single cells and would contribute small, but discernable variability differences to the total mRNA complement of a cell. They should exist in a defined chromosomal region and should generate mRNAs of a small set of adjacent genes within a specific open chromatin domain. Increased numbers of co-expressed genes arising from a wider chromosome region might suggest open chromatin regions that extend beyond typical Topologically Associated Domains (TADs) or Lamin Associated Domains (LADs)(5,6). This could be due to structural changes of the nuclei or incomplete polyploidy(7,8).

Here we demonstrate that simple statistical and data-driven methods can effectively evaluate the co-expression patterns of genes on the same chromosome within a single cell type. The outcome of this method is a full genome plot of the percent of co-expressed genes on a shared chromosome, where plot elevations indicate increased gene co-expression from the same chromosome. We termed the corresponding visualization a ‘Brooklyn plot’, due to similarities in appearance to the Manhattan plot of genome wide associations. We demonstrate that most normal (non-diseased) cells have a baseline of expected co-expression using scRNA-seq data, that the baseline is modestly increased in snRNA-seq data, and that rare diseased cell types can show a wide deviation from the non-diseased (baseline) pattern of co-expression in the same cell type. This deviation provides strong evidence of chromosomal domain dysregulation in a diseased cell setting.

## MATERIALS AND METHODS

### Establishing a search space

To enable investigation of chromosomal co-expression patterns, we developed the Brooklyn plot software package (https://github.com/arunhpatil/brooklyn) (Supplemental Figure 1). The input is an h5ad file of a scRNA-seq or snRNA-seq dataset obtained from any source. The h5ad datasets are loaded with the Scanpy package. An ‘observations’ list (h5ad.obs) is identified for the full h5ad dataset and the available criteria are used to filter down to one cell type with or without any additional user-selected delimiters such as disease state or tissue region. For these cells, raw gene expression is ranked and the 3500 most highly expressed genes becomes the full search space. This level of coverage provides for ∼1 gene per megabase (MB) throughout the genome. The Brooklyn plot package then obtains gene chromosomal locations from Biomart annotations and uses these to arrange the 3500 genes by chromosome location. Every 10^th^ gene is used as a standard marker for ∼10 MB coverage of the entire genome (n=350 genes). Alternatively, the user can individually select genes to serve as standard markers across the genome.

### Calculating chromosome co-expression

A Pearson correlation is made for each standard marker gene (n=350) to the full search space (n=3500) in an iterative fashion. For each standard marker gene, the correlation is ranked by a Bonferroni corrected p-value. The 50 highest correlated genes are then subselected. From this group, it is determined what percent of these genes are also present on the same chromosome. For example, if the standard marker gene was located on chromosome 3, and 10 of the 50 highest correlated genes were on chromosome 3, then the value is 20%. Increases of same-chromosome gene expression above chance, indicate larger co-transcriptional activities from the same chromosome. This approach (350/3500 genes) is computationally fast, while establishing a threshold in which elevations in broad chromosomal co-expression are easily detected. These data are used to generate a Brooklyn plot, based on the percent of chromosome co-expression for each of the 350 marker genes (Figure 1). Further detail of the methodological and analysis steps are described in Supplemental methods and documented as a Jupyter notebook (https://brooklyn-plot.readthedocs.io/).

**Figure 1.**
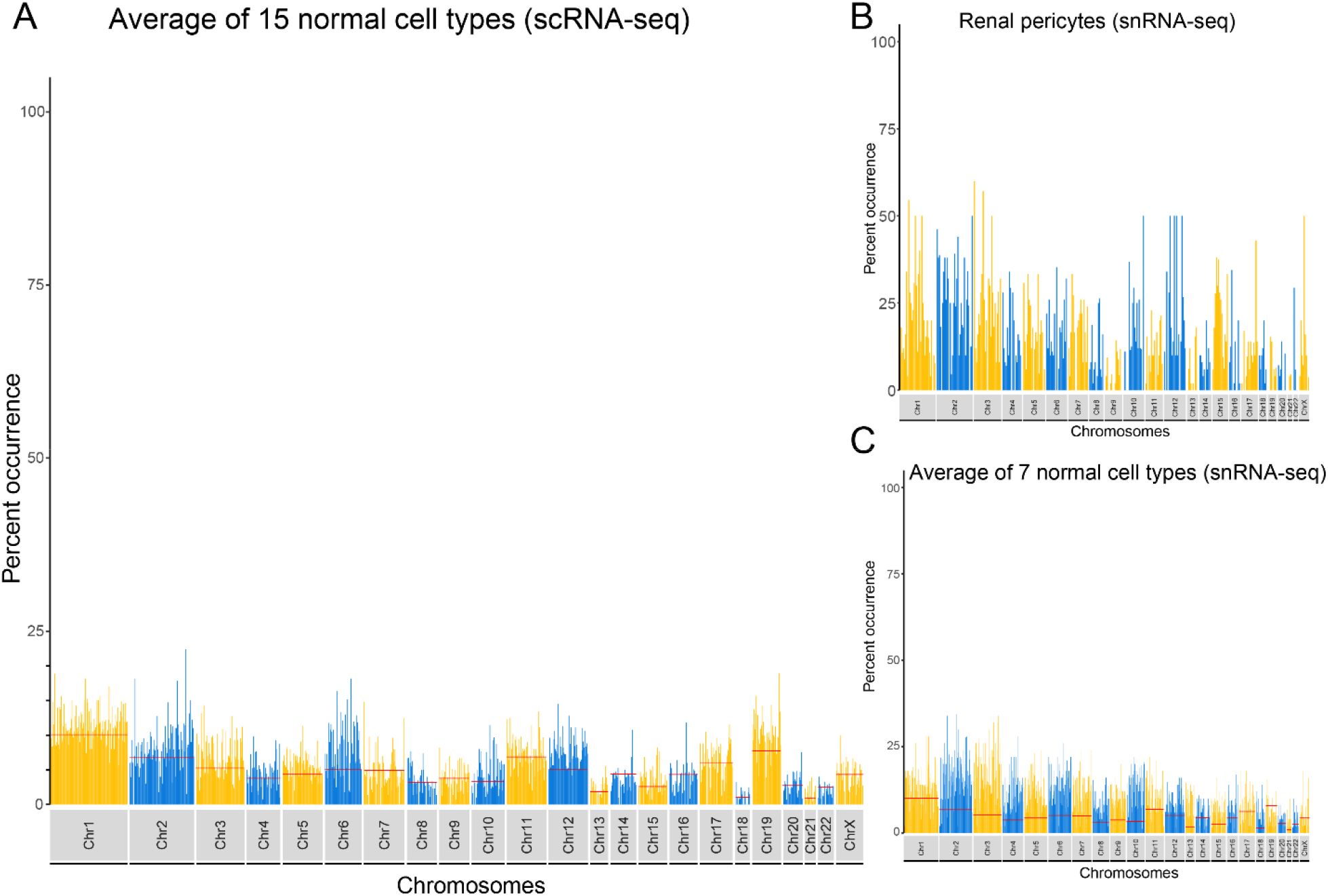
A) A combined Brooklyn plot from 15 non-diseased (normal) cell types obtained from 5 scRNA-seq studies. The percent occurrence is the percent of genes among the top 50 pairwise correlated genes, from the same chromosome. Red lines indicate values expected-by-chance per chromosome. B) Brooklyn plot of snRNA-seq data from 1962 renal pericyte cells, demonstrating modestly elevated (global 15.09%) co-expression of genes from the same chromosome across the genome. scRNA-seq results from matched renal pericytes was 5.89%. C) A combined Brooklyn plot from 7 non-diseased cell types obtained from two snRNA-seq studies.

## RESULTS

### Baseline level of chromosomal co-expression

Based on known patterns of gene expression on each chromosome (e.g. 9.6% of all genes are on chromosome 1 [Supplementary Table S1]), we established a baseline level of expected chromosome co-expression (Figure 1A, red bars). At a genome wide level, this is 4.35% although it varies by chromosome based on the number of genes on the given chromosome. We then used the Brooklyn plot package to obtain genome co-expression data from 40 experiments (17 cell types) across 6 normal-tissue and 6 diseased-tissue studies (Supplementary Table S2)(9-16). We combined the scRNA-seq data from 13 non-diseased cell types (15 samples, cell count ranges from 289 to 7205), for a baseline chromosome gene co-expression pattern (Figure 1A, Supplementary Methods). The average chromosomal co-expression for this group was 5.94%, which was slightly above the prior null expectation (4.35%), indicating a general lack of increased chromosomal co-expression.

### Co-expression in snRNA-seq and diseased cell types

For five matched cell types, snRNA-seq had double the amount of genome-wide co-expression to scRNA-seq (11.75% vs 5.93%, Figure 1B) and overall, snRNA-seq from 7 cell types had increased co-expression (Figure 1C). We observed very little difference in average co-expression values between normal and disease states for a variety of cells using either scRNA-seq or snRNA-seq datasets (Supplementary Table S2).

### Elevated co-expression in dilated cardiomyopathy

We then investigated a cell type in a disease state that we hypothesized might have an altered Brooklyn plot. Titin (TTN) is a structural protein of the cardiac muscle cell (cardiomyocyte) sarcomere. Mutations in *TTN* cause dilated cardiomyopathy and are associated with unusual/stellate nuclei(17). A Brooklyn plot of *TTN* mutated left ventricular cardiomyocytes showed marked global upregulation of chromosomal co-expression (61.14%, Figure 2A). Twenty-four standard genes had 49 or more of the top 50 correlated genes present on the same chromosome. These correlated genes were generally enriched around the standard gene of interest, with enrichment signal not crossing the p and q arms of the chromosome (Fig 2B). This indicated a wide, but structured (∼50 MB) region of co-transcription in the cells. A second dataset of genetically unclassified dilated cardiomyopathy validated these initial findings, with global values of 61.45% and 67.75% for two different cardiac muscle cell subsets (Supplementary Table S2)(15).

**Figure 2.**
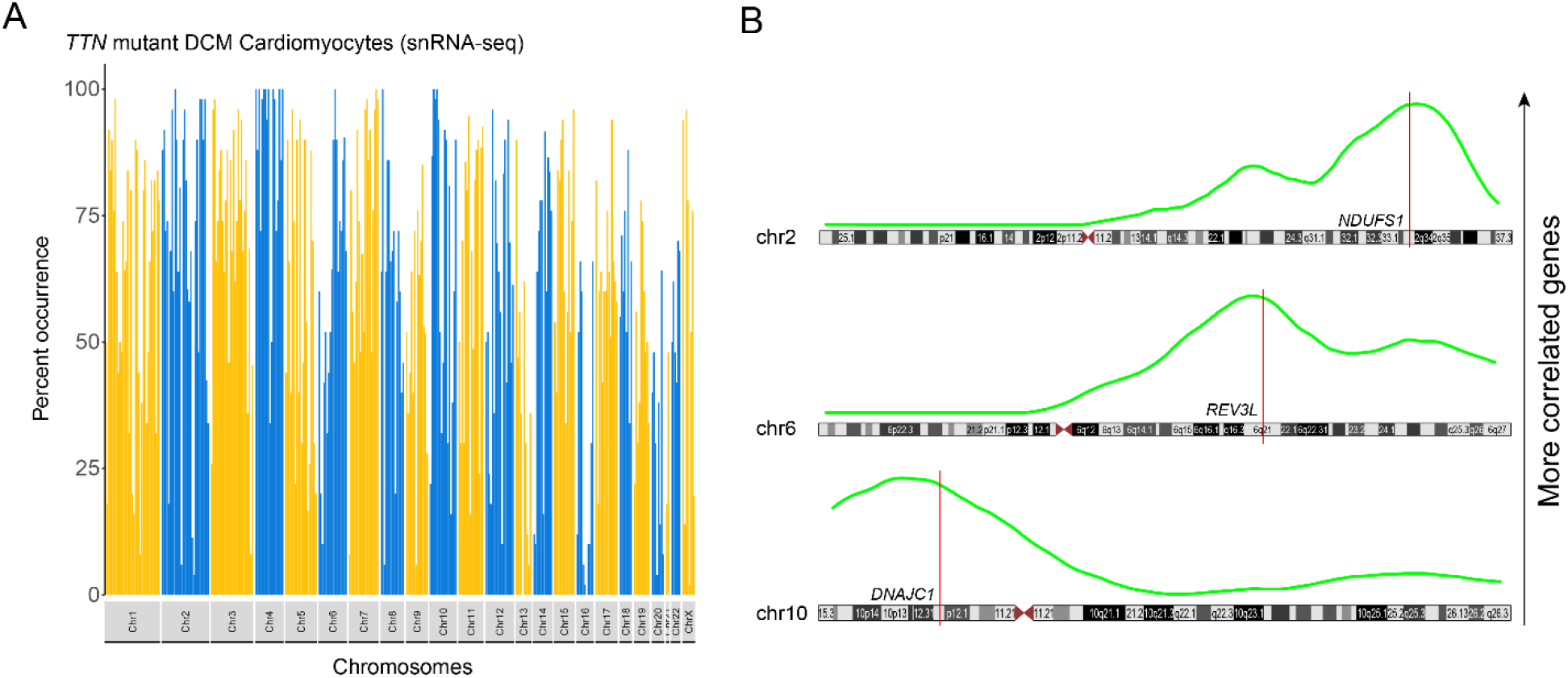
A) Brooklyn plot of *TTN* mutant DCM cardiomyocytes demonstrating marked elevation of co-expression (global 61.14%) suggesting altered open chromatin regions. B) Running average plots (green) for three standard genes (*NDUFS1, REV3L* and *DNAJC1*) in the *TTN* mutant DCM cardiomyocytes with ≥95% of the top co-expressed genes on the same chromosome. Peak values of co-expression localization are adjacent to the standard genes indicating increased neighboring co-expression rather than random distributions of co-expression, suggestive of larger open chromatin regions.

## DISCUSSION

Herein we describe a new method to evaluate aberrant chromosome co-transcription. We demonstrate that cell-to-cell variation within a well-defined cell cluster can uncover this previously unrecognized feature of extended co-transcription in single cell or single nuclei RNA-seq data. Consistent with prior expectations, there was no elevation of co-transcription for most cell types. This would be expected, as the Brooklyn plot measures elevations in a ∼50MB search space per standard marker gene, which is markedly larger than standard open chromatin domains. This further indicates that expression differences between cells in a defined cell cluster are features of other biologic (e.g. pathway activation) or technical (e.g. ambient RNA) causes(18).

We did observe increased global values for snRNA-seq samples compared to matched scRNA-seq samples. Nuclear mRNAs are generally the most recently transcribed mRNAs of a cell, containing more pre-mRNAs and introns (19). The doubling of average genome wide co-expression values in snRNA-seq over scRNA-seq, likely indicates more capture of transcriptional bursting in snRNA-seq data in which a small group of adjacent genes are transcribed at the same time.

The most important discovery using this method was establishing a marked increase in the co-transcription pattern in diseased cardiac muscle cells. Whereas normal cardiac muscle cells had values of 6.03%, diseased cells had 10x more chromosomal co-transcription. This finding is consistent with our established understanding of cardiac muscle cell nuclei. In the non-disease setting they are round to oval, but diseased cardiac muscle cell nuclei can be sharply angulated and non-ovoid (Figure 3A). As a result, we hypothesize that DNA becomes untethered from LADs attached to the nuclear envelope in a stochastic fashion between cardiac muscle cells, likely related to cell-specific mechanotransduction of intermediate filaments on the nuclear membrane (Figure 3B)(20). The altered nuclear structure likely alters open and closed (A/B) compartments allowing for transcription of genes to extend over ∼50 MB regions. This variability may also explain our previous discovery of 143 proteins with variable patterns of expression in these same cells(21). We entertained technical explanations for the elevated Brooklyn plots for cardiac muscle cells, but none could be found. Importantly, the data replicated across two separate datasets, but was not a feature of non-cardiac muscle cells from the same reports.

**Figure 3.**
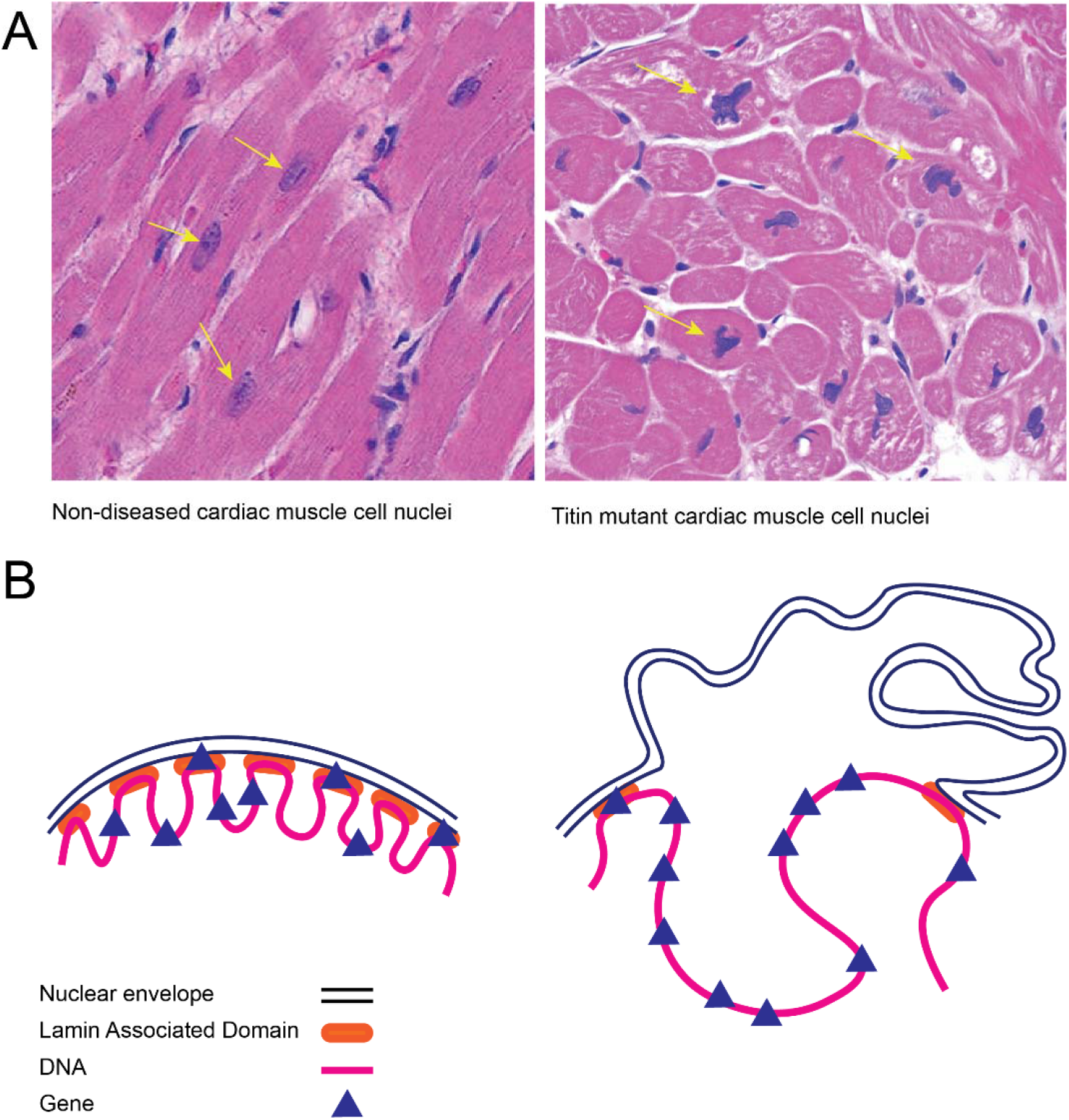
A) Images of cardiac muscle cell nuclei from non-diseased and *TTN* mutant hearts. The nuclei are smooth ovoid shapes in non-diseased cells (yellow arrows). In *TTN* mutant dilated cardiomyopathy, the nuclei are a variety of unusual and angulated shapes. B) A schematic of multiple Lamin Associated Domains (LADs), where the DNA is tethered creating separate areas of open/closed chromatin allowing few genes to be co-transcribed. To the right, with an irregular nuclear membrane, as seen in *TTN* mutant hearts, the LAD tethering is lost and larger regions of DNA are open to a transcriptional burst containing many more genes.

There are some key limitations to this method. The first is that scRNA-seq and snRNA-seq are sparse matrixes of data. Attempts to increase the search space from 3500 up to 7000 genes as a means to provide tighter co-expression windows around genes, did not improve the method due to expression sparsity. Secondly, the fully automated method of subselecting every 10^th^ gene from the 3500 gene window does not faithfully capture a gene every 10 MB, due to the known variability of gene rich/gene poor regions of the genome. Thus, while the average is ∼10 MB per gene, this can be variable and a source of randomness in the system. A time consuming, bespoke approach to identify 350 genes at 10 MB intervals was compared to this high throughput approach with negligible differences. Finally, attention is needed to make sure all cells of a named group within an h5ad file are within the same cluster. In one test of the method, a defined cell type in a public h5ad dataset had a subset of the cells intermixed in other clusters away from the primary cluster. The genes driving these variable locations featured consistently in all Pearson correlation datasets negating the value of the method. Thus, a solitary cluster of the participating cells, should be ensured before initiating the Brooklyn plot method.

In summary, we have developed a new package to generate Brooklyn plots, which can be used to explore altered chromosomal gene co-expression patterns indicating large-scale alterations in chromosomal regulatory domains. We expect as more datasets are generated and evaluated, that Brooklyn plots will uncover more transcriptional dysregulation in cells types across a range of diseases including malignancies.

## DATA AVAILABILITY

All h5ad datasets were obtained from the CellxGene portal (https://cellxgene.cziscience.com/) or the Single Cell Portal (https://singlecell.broadinstitute.org/single_cell) as indicated in Supplementary Data Table 2. The Brooklyn plot package source code is available at GitHub (https://github.com/arunhpatil/brooklyn).

## ACKNOWLEDGEMENTS

We thank Dan Arking, Aravinda Chakravarti, Loyal Goff, and Andrew McCallion for their helpful discussions.

---

**Supplementary Figure 1.**
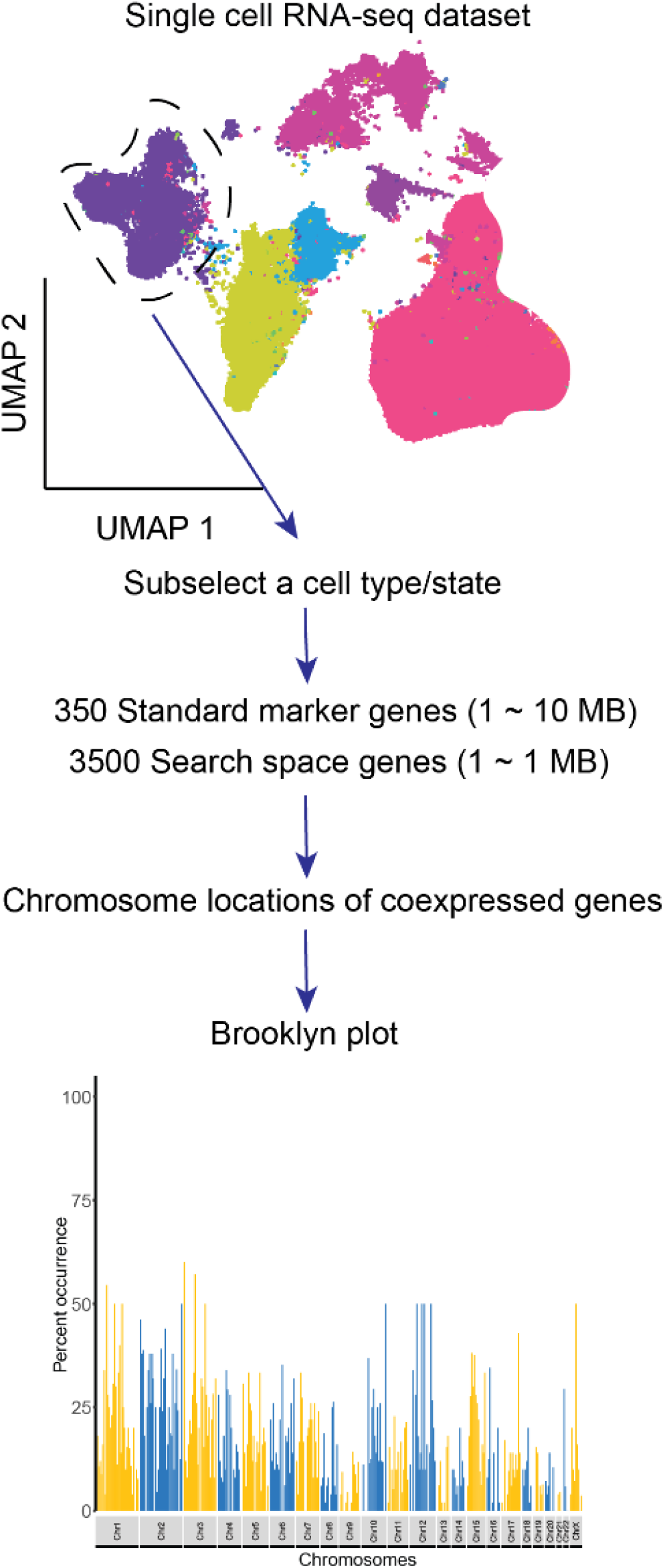
Schematic of the Brooklyn plot method. From a h5ad file, a single cell type/state is subselected from a scRNA-seq or snRNA-seq dataset (dotted circle in the UMAP plot). Standard marker genes and a search space are established. Co-expression by chromosome is established and a Brooklyn plot is generated.

**Supplementary Table 1**. The percent of genes across each chromosome. Data from Ensembl.

**Supplementary Table 2**. Metadata on 40 samples used in this study.

**Supplementary Methods**. Methods for using Brooklyn plot and specific methods for methods specific to this manuscript.

