## Supplementary Methods for "Brooklyn plots to identify co-expression dysregulation in single cell sequencing"

**Supplementary Methods Brooklyn plot functionality**

The Brooklyn plot package is available at GitHub (<https://github.com/arunhpatil/brooklyn>). It can be installed using conda “conda install -c bioconda brooklyn_plot” or PyPi (“python3.8 -m pip install --user brooklyn_plot”)(1). The input is an h5ad file of a single cell/nucleus sequencing dataset obtained from any source. If the single cell dataset of interest is not in h5ad format, it can be converted through sceasy (<https://github.com/cellgeni/sceasy>)(2).

To demonstrate the usefulness of the Brooklyn plot, we provide the h5ad file “Cardiomyocytes” from Cellxgene (https://cellxgene.cziscience.com/)(3). The dataset is loaded with the Scanpy package and is filtered to the user’s interests. In this example, only cardiomyocytes from left ventricles of patients with *TTN* mutant dilated cardiomyopathy were subselected. Other filters can be chosen with the goal to settle on a single cell type population. For these subsetted cells, raw gene expression abundance is ranked and the top 3500 expressed genes becomes the full search space, discarding more lowly expressed genes. Gene chromosomal locations are obtained from Biomart annotations. These genes are sorted by chromosome location and every 10^th^ gene is used as a standard for ~10 MB coverage of the entire genome (n=350 genes). The detailed analysis steps are documented as a Jupyter notebook (GitHub link). Output files are a subsetted h5ad file, the Biomart annotations, a list of 350 standard (to be queried) genes, and the 3500 genes in the search space.

These files are provided as input to the second script “brooklyn_plot,” where an internal module “brooklyn_arch()” converts the h5ad to a Numpy array and an iterative Pearson correlation for each query gene across the search space is performed using the “stats.pearsonr” method of Scipy. This generates a folder of CSV files for each gene that contains correlation coefficient (*r*), P-value and Bonferroni corrected P-value, -log_10_(P-value) and gene coordinates for all correlated genes above a Bonferroni corrected P-value of 0.05. The “summarize()” function in the Brooklyn plot package, determines the percent of the top 50 co-expressed genes (to all 350 standard genes) that localize to the same chromosome as the queried standard gene. Subsequently, this resultant summary file is represented as a Brooklyn plot in PDF format using the “ggplot2” package in R. The Brooklyn plot package utilizes “concurrent.futures” functionality and offers parallel processing across multiple threads for productivity gains.

**Combined Brooklyn plots**

For the 15-cell type scRNA-seq and 7-cell type snRNA-seq samples, the individual standard genes varied between samples. To overcome this, data was merged based on chromosome location in 1 MB intervals across the genome. Then all 1 MB intervals with a denominator of <50 genes were removed and the remaining values (percents of co-expressed genes from the same chromosome) were plotted. For the 15-cell type figure, this was 1065 measures. For the 7-cell type figure, this was 1083 measures.

**Localization of peak co-expression**

Three standard genes (*NDUFS1, REV3L, DNAJC1*) were selected from different chromosome locations from the *TTN* DCM output files. They were chosen for each having 98+% of the top 50 co-expressed genes being on the same chromosome. The location of the 49 or 50 co-expressed genes were solved relative to all genes on the chromosome (within the 3500 gene search space). A moving sum of the co-expressed genes for 15 adjacent genes was determined across the chromosome. Values ranged from 0 to 13. Then a moving average for 30 adjacent counts was used to smooth the values for plotting, which was performed in R.
